# Identifying functional multi-host shuttle plasmids to advance synthetic biology applications in *Mesorhizobium* and *Bradyrhizobium*

**DOI:** 10.1101/2023.12.20.572671

**Authors:** Jordyn S. Meaney, Aakanx K. Panchal, Aiden J. Wilcox, George C. diCenzo, Bogumil J. Karas

## Abstract

Ammonia availability has a crucial role in agriculture as it ensures healthy plant growth and increased crop yields. Since diazotrophs are the only organisms capable of reducing dinitrogen to ammonium, they have a great ecological importance and potential to mitigate the environmental and economic costs of synthetic fertilizer use. Rhizobia are especially valuable being that they can engage in nitrogen-fixing symbiotic relationships with legumes, and they demonstrate great diversity and plasticity in genomic and phenotypic traits. However, few rhizobial species have improved genetic tractability for synthetic biology applications. This study established a basic genetic toolbox with antibiotic resistance markers, multi-host shuttle plasmids and a streamlined protocol for biparental conjugation with *Mesorhizobium* and *Bradyrhizobium* species. We identified two *repABC* origins of replication from *Sinorhizobium meliloti* (pSymB) and *Rhizobium etli* (p42d) that were stable across all three strains of interest. Furthermore, the NZP2235 genome was sequenced and phylogenetic analysis determined its reclassification to *Mesorhizobium huakuii*. These tools will enable the use of plasmid-based strategies for more advanced genetic engineering projects and ultimately contribute towards the development of more sustainable agriculture practices by means of novel nitrogen-fixing organelles, elite bioinoculants or symbiotic association with non-legumes.

**GRAPHICAL ABSTRACT:** 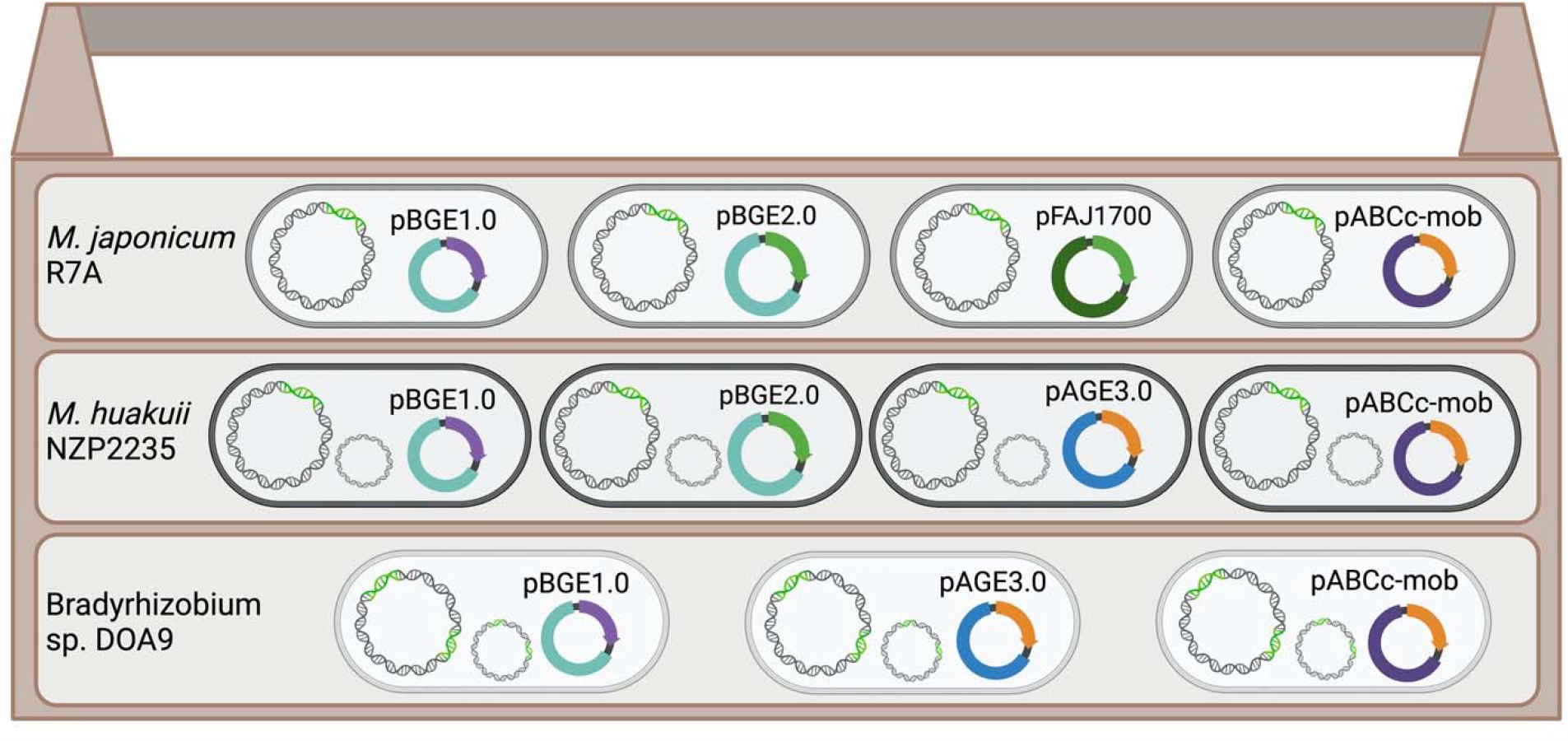

## INTRODUCTION

Nitrogen has a fundamental role in all biological systems, as it is a component of genetic material and proteins that govern cellular operations. However, dinitrogen, the most abundant form in nature, is biologically inactive due to the inert triple bond. This leaves autotrophic organisms, like plants, to rely on limited reserves of nitrates and ammonium found in soil and water, and whose low abundance often limits crop yields. The resulting bottleneck, known as the nitrogen problem, has been the most profound agricultural issue of the 21^st^ century. Application of synthetic fertilizers has become widespread, but this temporary solution is economically and ecologically costly, and highlights the need for more advanced engineering strategies that create a permanent supply of fixed nitrogen in crop varieties.

Biological nitrogen fixation (BNF) refers to the enzyme catalyzed reaction that converts atmospheric dinitrogen gas into bio-available ammonia. BNF is limited to a subset of prokaryotes, called diazotrophs, which harbour highly conserved nitrogen-fixation (*nif*) clusters encoding nitrogenase - a two component metalloenzyme – as well as proteins for cofactor assembly and electron transfer (Dixon and Kahn, 2004). This valuable mechanism is utilized by some diazotrophic bacteria in mutualistic associations, known as symbiotic nitrogen fixation (SNF), but requires expression of additional symbiotic nodulation (*nod/nol/noe*) genes (Wang, Liu and Zhu, 2018). The order *Hyphomicrobiales* encompasses 38 gram-negative bacterial families (List of Prokaryotic names with Standing in Nomenclature; LPSN; accessed 2 December 2023), including seven families (*Rhizobiaceae, Phyllobacteriaceae, Brucellaceae, Nitrobacteriaceae, Methylobacteriaceae, Xanthobacteriaceae*, and *Devosiaceae*) containing at least one rhizobial species. Through a complex cooperative infection process, these soil microbes can take up residence in root nodules of legume species, where photosynthates are supplied from the plant to internalized and differentiated bacteria (known as bacteroids) in exchange for fixed nitrogen in the form of ammonium (Udvardi and Poole, 2013). Amongst rhizobia there is great inter- and intra-species phenotypic and genomic diversity that influences adaptation to various soil environments, host specificity, and symbiotic efficiency (diCenzo *et al*., 2019; Geddes *et al*., 2020; Walker, Lagunas and Gifford, 2020).

Progress towards creating self-fertilizing plants has focused on two main areas: transferring the genetic components necessary for BNF to plant hosts (Li and Chen, 2020) and expansion of symbiotic compatibility for SNF between rhizobia and non-legume hosts (Guo *et al*., 2023). More recently, a proposal to develop novel nitrogen-fixing organelles from free-living diazotrophic bacteria using principles of endosymbiosis was presented as an alternative strategy (Meaney *et al*., 2020). Utilizing whole bacteria circumvents the need to refactor *nif* clusters (Temme, Zhao and Voigt, 2012), provides a microaerobic environment that supports oxygen-labile nitrogenase activity (Burén and Rubio, 2018), and creates a compartment for nitrogen fixation separate from oxidative phosphorylation and photosynthesis. Not only do rhizobia perform the desired function, but their propensity to participate in intracellular symbiotic relationships makes them ideal candidates as primitive organelles. Furthermore, the naturally dynamic lifestyle and genomic plasticity displayed by rhizobia may aid in the transformation from free-living organism to organelle (Wang *et al*., 2020; Hawkins and Oresnik, 2022).

Of the fast-growing rhizobia, *Sinorhizobium meliloti* has been heavily characterized and a wide array of genetic tools - including multi-host shuttle plasmids, metabolic modeling, and DNA editing techniques - as well as reduced and minimal N_2_-fixing genomes have been developed in recent years (Döhlemann *et al*., 2017; diCenzo *et al*., 2018, 2019, 2020; Brumwell *et al*., 2019). Meanwhile, intermediate-(*Mesorhizobium*) and slow-growing (*Bradyrhizobium*) rhizobia have received less attention towards advancing their genetic tractability (diCenzo *et al*., 2019). These families generally have their SNF genes (*nif, fix*, and *nod*) organized in a chromosomal symbiosis island, unlike *Rhizobium* and *Sinorhizobium* strains, which feature a symbiotic megaplasmid (Figure 1A) (Sullivan, Brown and Ronson, 2013; Geddes *et al*., 2020). The model *Mesorhizobium* strain, *M. japonicum* MAFF303099, has a multipartite genome organization, while the closely related strain *M. japonicum* R7A has a single chromosome (Kaneko *et al*., 2000; Kelly *et al*., 2014). Additionally, *M. japonicum* exhibits a broad legume host range, whereas some other mesorhizobia, as is typical of Mesorhizobium *huakuii* strains, have a narrow host range. Within the family *Bradyrhizobiaceae*, one broad host range strain that has been well characterized in recent years is of particular interest to us (Teamtisong *et al*., 2014; Songwattana *et al*., 2017; Wongdee *et al*., 2023). *Bradyrhizobium* sp. DOA9, first described in 2012 (Noisangiam *et al*., 2012), exhibits differential expression of two distinct sets of chromosomal and plasmid encoded *nif* genes, which accommodates nitrogen-fixation in symbiotic or free-living states (Figure 1B) (Wongdee *et al*., 2016, 2018). Harnessing this unique feature for the development of a nitrogen-fixing organelle could greatly reduce the anticipated difficulties in triggering symbiotic activation of the pathway in non-legumes.

**Figure 1.**
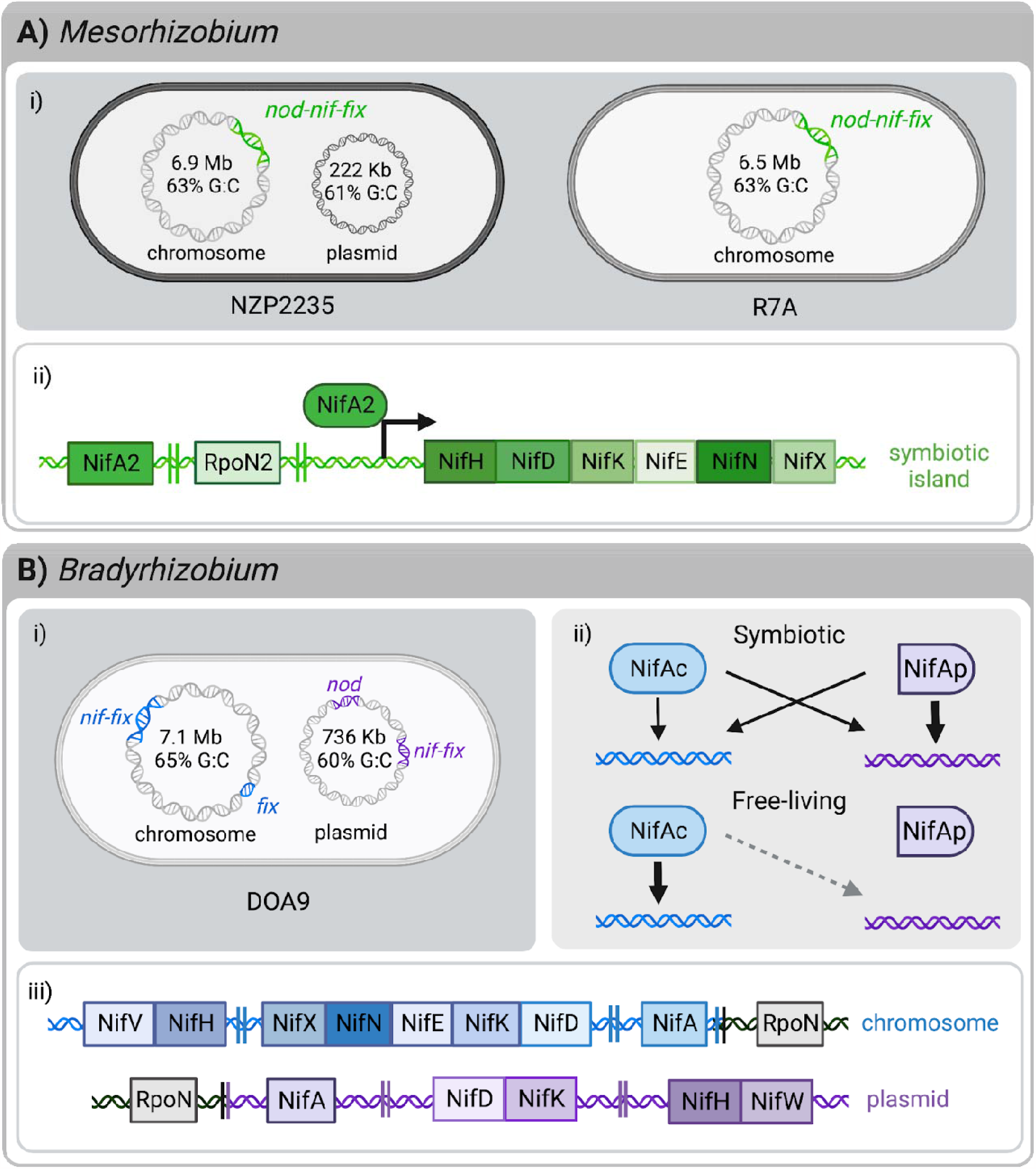
Comparative overview of characteristics. **(A)** *Mesorhizobum huakuii* NZP2235 and *Mesorhizobium japonicum* R7A. i) NZP2235 has a 7.3 Mb genome with a chromosome and plasmid, while R7A has a single chromosome of 6.5 Mb. Both species have a 502-kb symbiosis island (ICESym) on the chromosome. ii) Regulation of nitrogen-fixation genes in mesorhizobia is dependent on expression of *rpoN2* and *nifA2*. Structural genes for nitrogenase are located adjacent to each other in a single operon. **(B)** *Bradyrhizobium* sp. DOA9. i) The 7.8 Mb genome is composed of a chromosome and a plasmid, both of which have two smaller regions encoding nodulation or nitrogen fixation genes. ii) Nitrogen fixation regulation is distinct in symbiotic and free-living growth. During symbiosis, *nifAc* and *nifAp* are redundant, but *nifDKp* is primarily responsible for nitrogenase activity. In the free-living state, *nifAc* is indispensable and *nifDKc* is the major contributor to nitrogenase expression. iii) Nitrogen-fixation genes are present on both the chromosome and plasmid. In both cases, the structural genes for nitrogenase are split into two operons. Accessory protein *nifV* is also located on the chromosome which may contribute to nitrogen-fixation in the free-living state.

Engineering endosymbionts will be essential to the establishment and maintenance of a stable relationship with the host in early stages of novel organelle evolution, as demonstrated by Mehta and Cournoyer (Mehta *et al*., 2018, 2019; Cournoyer *et al*., 2022). Considering the variability across rhizobia species, there is value in examining the viability of strains with different genomic and phenotypic backgrounds in directed endosymbiosis (Meaney *et al*., 2020). This study intends to characterize functional selection markers and replication origins for propagation of multi-host shuttle plasmids as well as a mechanism for DNA delivery to *M. japonicum* R7A, *M. huakuii* NZP2235, and *Bradyrhizobium* sp. DOA9. Identification of basic genetic tools will enable plasmid-based engineering strategies that facilitate more advanced synthetic biology techniques (Nora *et al*., 2019). Aside from the development of novel organelles, these organisms could also be manipulated for other applications, including the development of elite bioinoculants or expanding symbiosis to non-legumes (Checcucci *et al*., 2017; Pankievicz *et al*., 2019; Krönauer and Radutoiu, 2021; Benmrid *et al*., 2023).

## MATERIALS AND METHODS

### Strains and culture conditions

All bacterial strains used in this study are listed in Table S1 with the appropriate antibiotic concentrations. *Escherichia coli* strains were grown in Luria Broth (LB) (10 g/L tryptone, 5 g/L yeast extract, 10 g/L NaCl) at 37°C. All rhizobia were grown at 30°C. *Bradyrhizobium* sp. DOA9 was grown in YEM (1 g/L yeast extract, 10 g/L D-mannitol, 0.5 g/L K_2_HPO_4_, 0.2 g/L Mg_2_SO_4_, and 0.1 g/L NaCl, and adjusted to pH 7.0). *M. huakuii* NZP2235 and *M. japonicum* R7A were grown in modified TY (TY-M) (5 g/L tryptone, 3 g/L yeast extract, 1 g/L D-mannitol and 0.66 g/L CaCl_2_ and adjusted to pH 7.0).

### Streamlined conjugation method

*Preparation of* E. coli. The donor *E. coli* ECGE101 strains were streaked on LB agar plates from glycerol stock with diaminopimelic acid (60 µg/mL) and the appropriate antibiotics, before being incubated at 37°C degrees overnight. A single colony for each *E. coli* strain was used to inoculate 3 mL starter cultures shaking at 225 rpm at 37°C. Cultures were diluted 50x into fresh media and grown to an OD_600_ of 0.6; if cultures exceeded the target density, they were adjusted back to an OD_600_ of 0.6 in 50 mL. Cells were collected by centrifugation for 15 minutes at 3,000 x *g*. The supernatant was decanted and all residual liquid was removed. Pellets were resuspended in 1 mL of 10% glycerol. After transferring 125 µL aliquots to 1.5 mL Eppendorf tubes, the cells were flash frozen in -80°C ethanol and stored at -80°C.

#### Preparation of rhizobia

Wildtype recipient strains were streaked on non-selective media from glycerol stock and incubated at 30°C for 4-8 days for single colonies to emerge. *Mesorhizobium* single colonies were streaked onto fresh media and incubated for two days. *Bradyrhizobium* single colonies were spread across a 1 x 3 cm patch and incubated for 4 days. For inoculation of 50 mL cultures, a pipette tip worth of cells was used for *Mesorhizobium* while the entire streak of cells was used for *Bradyrhizobium*. The cultures were incubated at 30°C, shaking at 225 rpm, and grown for one or two days for *Mesorhizobium* and *Bradyrhizobium*, respectively. If cultures exceed the target density, they were adjusted back to an OD_600_ of 0.6 in 50 mL. Cells were collected by centrifugation for 15 minutes at 3,000 x *g* at 4°C. The supernatant was decanted and all residual liquid was removed. Pellets were resuspended in 1 mL of 10% glycerol. After transferring 250 µL aliquots to 1.5 mL Eppendorf tubes, the cells were flash frozen in -80°C ethanol and stored at -80°C. See Figure S1A.

#### Conjugation

Frozen aliquots were removed from the -80°C and thawed on ice for 30 minutes. Once thawed, cells were vortexed before transferring 100 µL aliquots of each recipient and donor strain onto agar plates supplemented with diaminopimelic acid (60 µg/mL). Controls contained 100 µL of sterile water with the donor or recipient strain. Cells were spread evenly, and plates were allowed to dry before incubation at 30°C for 3 hours. After conjugation, the plates were scraped with 1 mL of sterile water and the cell suspension was transferred to an Eppendorf tube. The volume of the cell suspension was adjusted to a total volume of 1 mL with sterile water, as needed. Suspensions were used for a 10x dilution series (10^-1^ to 10^-3^) and 150 µL of each dilution was spread on recovery plates with the appropriate antibiotic selection (Table S1). Plates were incubated at 30°C for 3-7 days for single colonies to form. Colonies were passed on selective media twice prior to screening. See Figure S1B.

#### Antibiotic Spot Assays

Frozen 250 µL aliquots of rhizobia, prepared as described above, were defrosted on ice for 30 minutes. Cells were transferred to 50 mL of fresh media and incubated at 30°C with shaking at 225 rpm for 3 hours. Cultures were then centrifuged for 15 minutes at 3,000 x *g* at 4°C. Supernatant was removed and the samples were then centrifuged again using the same conditions for 10 minutes. Residual culture was removed and the pellet was resuspended in 500 µL of sterile water. The resuspensions were adjusted to an OD_600_ of 6.0 and 10-fold dilutions were performed. Spots were plated on a range of antibiotics of varying concentrations and on non-selective media as a control. Each spot consisted of 4 µL of the respective dilution from 10^-1^ to 10^-6^. Spot assays were performed in triplicate with single wildtype colonies. Plates were incubated for two days for *Mesorhizobium* and four days for *Bradyrhizobium* before pictures were captured.

#### Whole Genome Sequencing

A single colony of *M. huakuii* NZP2235 was inoculated into TY-M broth and grown until saturation, after which DNA was extracted using Monarch genomic DNA purification kit (New England Biolabs) according to the manufacturer’s instructions. Oxford Nanopore Technologies (ONT) sequencing and basecalling with Guppy 6.4.6+ae70e8f, as well as Illumina sequencing, were performed as described elsewhere (Kaur *et al*., 2023). The Illumina reads were filtered using BBDuk 38.96 and trimmed using Trimmomatic 0.39 (Kaur *et al*., 2023). Genome assembly, polishing, and annotation were performed as described previously, using the following software: Flye 2.9-b1779, Medaka 1.7.2, polypolish 0.5.0, POLCA 4.0.9, bwa 0.7.17-r1198-dirty, Circlator 1.5.5, and PGAP 2023-10-03.build7061 (Kaur *et al*., 2023). All sequencing, assembly, and annotation data can be accessed via NCBI BioProject PRJNA1044807.

#### Phylogenetic Analyses

The Type (Strain) Genome Server (Meier-Kolthoff and Göker, 2019; Meier-Kolthoff *et al*., 2022) was used to identify *Mesorhizobium* and *Bradyrhizobium* species type strains related to *M. japonicum* R7A, *M. huakuii* NZP2235, and *Bradyrhizobium* sp. DOA9. Genome sequences for the identified type strains were then downloaded from the National Center for Biotechnology Information (NCB) Genome database. ANI was calculated between each pair of genomes using FastANI 1.33 (Jain *et al*., 2018). GET_HOMOLOGUES 05052023 was used with custom scripts to identify single-copy marker genes present in 100% of the target genomes, as described previously (diCenzo *et al*., 2023). The encoded proteins were then used to construct a phylogenomic tree as described previously using the following software: MAFFT 7.453, trimAl 1.4.rev22, and IQ-TREE 2.2.2.4 (diCenzo *et al*., 2023). Phylogenies were visualized using iTOL (diCenzo *et al*., 2023).

#### DNA Isolation for Screening

Plasmid DNA from *E. coli* was obtained using the BioBasic EZ-10 miniprep kit, according to the manufacturer’s instructions, with 3 mL of saturated overnight culture. Strains carrying pAGE/pBGE replicons were induced with arabinose (100 µg/mL). Alkaline lysis was performed for DNA isolation from all rhizobia using 5-10 mL cultures, grown for 2 or 4 days for *Mesorhizobium* and *Bradyrhizobium*, respectively. Please refer to the supplementary methods for the alkaline lysis procedure.

#### Multiplex PCR Screening

Transconjugants were screened separately for plasmid and genomic DNA using the Qiagen Multiplex PCR kit with the primers listed in Table S4, according to manufacturer’s instructions. The “Standard Multiplex PCR Protocol” was followed, adjusting the components for a final volume of 20 µL with 1 µL of undiluted DNA isolated using alkaline lysis as described above. PCRs were performed using a 3-step PCR cycling protocol consisting of: a melting temperature of 94°C for 30 seconds, an annealing temperature of 60°C for 90 seconds, and an extension temperature of 72°C for 45 seconds, repeated 35 times. Gel electrophoresis was performed with 1 µL of multiplex PCR (MPX) product run on a 2% agarose gel.

## RESULTS

### Whole genome sequence of M. huakuii NZP2235

Since the genome of the rhizobial strain NZP2235 was not yet available online, high molecular weight DNA was isolated and sequenced. Phylogenomic and ANI data indicated that NZP2235 is most closely related to *M. huakuii* NBRC 15243^T^ (Figure 2) and thus NZP2235 was reclassified as *M. huakuii* NZP2235 (formerly *M. loti* NZP2235 (Jarvis, Pankhurst and Patel, 1982; Karas *et al*., 2005). The assembly revealed a multipartite genome split between two replicons, a chromosome (GenBank accession CP139858) and a plasmid (pMhuNZP2235a; CP139859), with 6,990,725 bp (63.1 G:C) and 221,974 bp (60.8% G:C), respectively (Figure 1A). Like many endogenous rhizobial plasmids, pMhnNZP2235a encodes *repABC* family proteins for replication initiation and propagation as well as mating pair formation genes involved in conjugative mobilization (diCenzo and Finan, 2017; Geddes *et al*., 2020). However, it does not contain symbiotic genes, indicating that it may instead contribute to competition in the soil or for root nodule occupancy. Instead, the symbiotic island was found on the chromosome following the *tRNA-Phe* gene and shared 99.9% pairwise identity with the integrative and conjugative element (ICESym) of *M. japonicum* R7A (Figure S2) (Colombi *et al*., 2021).

**Figure 2.**
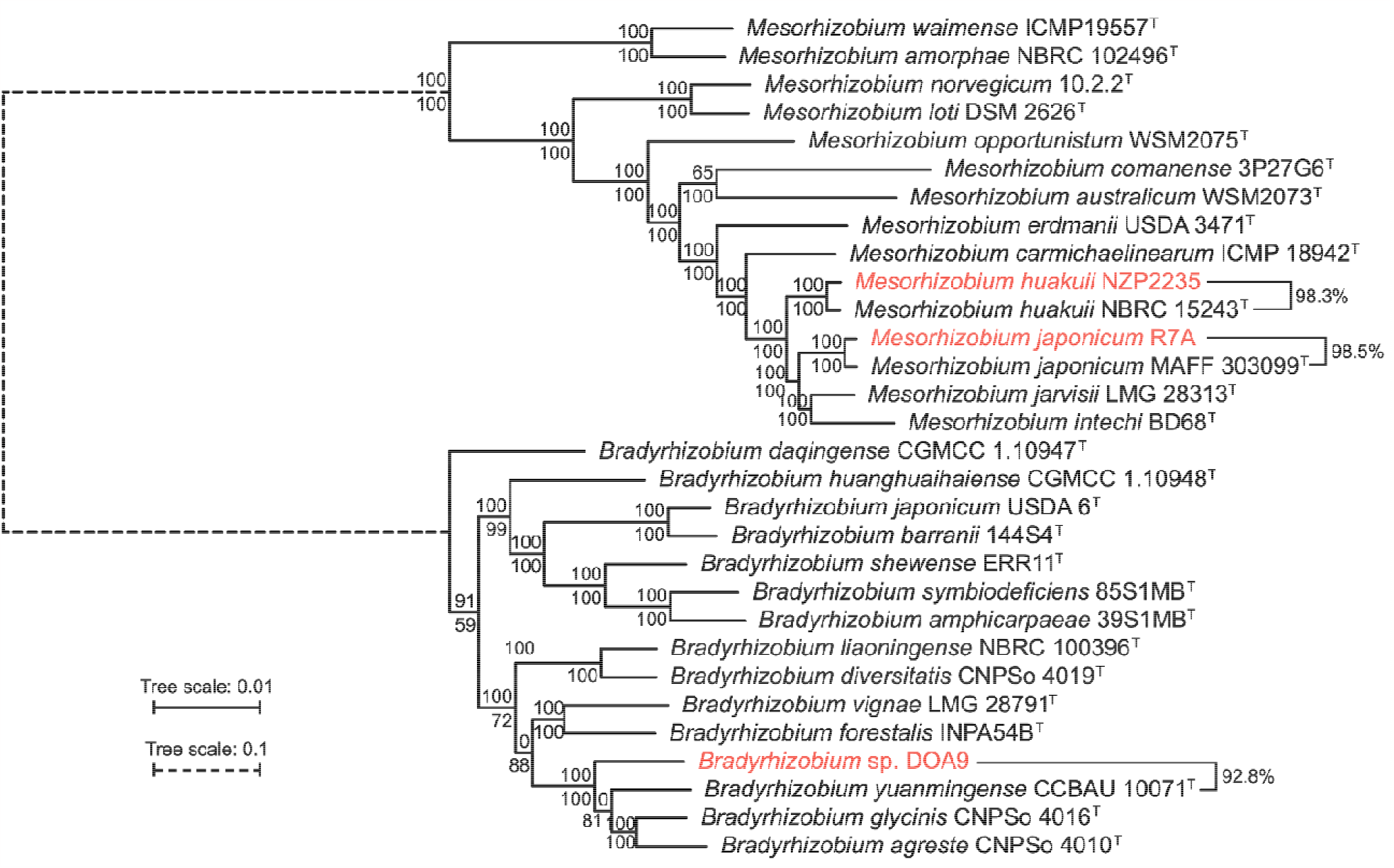
Maximum likelihood phylogeny of the genera *Mesorhizobium* and *Bradyrhizobium*. A maximum likelihood phylogeny of the three focal strains (shown in red) together with 27 species type strains, constructed from the concatenated protein alignments of 1,168 core proteins. The numbers on the nodes indicate the Shimodaira-Hasegawa-like approximate likelihood ratio test (SH-aLRT) support values (top numbers) and the ultra-fast jackknife values using a 40% resampling rate support values (bottom numbers), both calculated from 1000 replicates. The scale bars represent the average number of amino acid substitutions per site; for presentation purposes, a different scale was used for the branch connecting the two genera. Average nucleotide identity (ANI) values are given on the right side of the figure for comparisons between the three focal strains and the most closely related species type strains.

### Assessing antibiotic sensitivity

The response to a variety of antibiotics was evaluated using spot assays for each wildtype strain on media supplemented with a range of concentrations for each compound (Figures S3-5). Growth was compared against non-selective media to assess sensitivity to each antibiotic. *M. huakuii* NZP2235 displayed moderate resistance to ampicillin and streptomycin, and high resistance to kanamycin. Similarly, *M. japonicum* R7A showed moderate resistance to ampicillin and kanamycin, but high concentrations were successful at suppressing growth. *Bradyrhizobium* sp. DOA9 was the least sensitive to antibiotic exposure, and was highly resistant to chloramphenicol, gentamicin, and kanamycin, rendering genes providing resistance to these antibiotics ineffective as marker genes for positive selection in this strain. Based on spot assays, the minimum concentrations for selection with each antibiotic are listed in Table 1 for all three organisms. Selection at these concentrations, or higher, after DNA delivery will ensure that untransformed cells do not survive.

**Table 1.** Suggested concentrations for antibiotic selection based on natural sensitivity. Colour indicates the strength of antibiotic selection (green – good; yellow – weak; red – not effective). All concentrations are reported in µg/mL.

| Antibiotic | NZP2235 | R7A | DOA9 |
| --- | --- | --- | --- |
| Ampicillin | 250 | 150 | 250 |
| Chloramphenicol | 15 | 20 | N/A |
| Gentamycin | 40 | 20 | N/A |
| Kanamycin | N/A | 250 | N/A |
| Neomycin | 100 | 30 | 150 |
| Spectinomycin | 30 | 30 | 100 |
| Streptomycin | 250 | 100 | 50 |
| Tetracycline | 5 | 1 | 50 |

### Streamlined conjugation protocol

Conjugation was the preferred DNA delivery mechanism since wildtype rhizobia are known to encode restriction-modification (R-M) systems that reduce the efficiency of traditional transformation methods, like heat shock and electroporation (Ferri *et al*., 2010; Riley and Guss, 2021). Double stranded plasmid DNA used in transformation is isolated with a host-specific methylation pattern that is recognized by recipient methylases as foreign information and subsequently targeted for degradation (Murray, 2000). Conversely, during bacterial conjugation, a single strand of plasmid DNA is transferred across the host pilus into the recipient, where replication is completed (Waksman, 2019). The resulting hemi-methylated DNA prevents effective catalytic activity by restriction endonucleases, facilitating evasion of the R-M system (Murray, 2000; Bochtler, 2021). However, it has been reported that intact R-M systems may still cause a deficit in conjugation efficiency (Roer, Aarestrup and Hasman, 2015). Currently R-M systems are unmapped on rebase.neb.com for all three strains, but several genes of interest have been identified (Table S2), which could be targeted for disruption in future studies using plasmid-based editing techniques.

Triparental mating has been the most common strategy for conjugative plasmid transfer in rhizobia, utilizing a helper, donor, and recipient strain. Instead, the method used here employs only two strains by including a mobilization plasmid, pTA-Mob, as well as the conjugative shuttle plasmid in the *E. coli* donor (Strand *et al*., 2014). Moreover, absence of *dapA* in the donor strain (*E. coli* ECGE101) renders it a diaminopimelic acid auxotroph and allows wildtype rhizobial transconjugants to be easily obtained through counterselection by withholding DAP supplementation in the selection media (Brumwell *et al*., 2019). For convenience, a segmented procedure was created to separate bacterial culturing from conjugation as it is laborious to coordinate harvest of fast-growing *E. coli* with the slow-growing recipient strains (Figure 3). All bacteria were grown to exponential phase to ensure the harvest of healthy cells prior to flash freezing. Donor *E. coli* can be prepared in four days from frozen stock, while *Mesorhizobium* and *Bradyrhizobium* recipient strains require 8 or 14 days, respectively (Figure S1). Prepared cells can be stored indefinitely, and multiple replicates of an experiment can be run independently from the same batch. The streamlined protocol is further simplified by avoiding the use of nitrocellulose filters and having a shortened three-hour conjugation period compared to the traditional overnight incubation.

**Figure 3.**
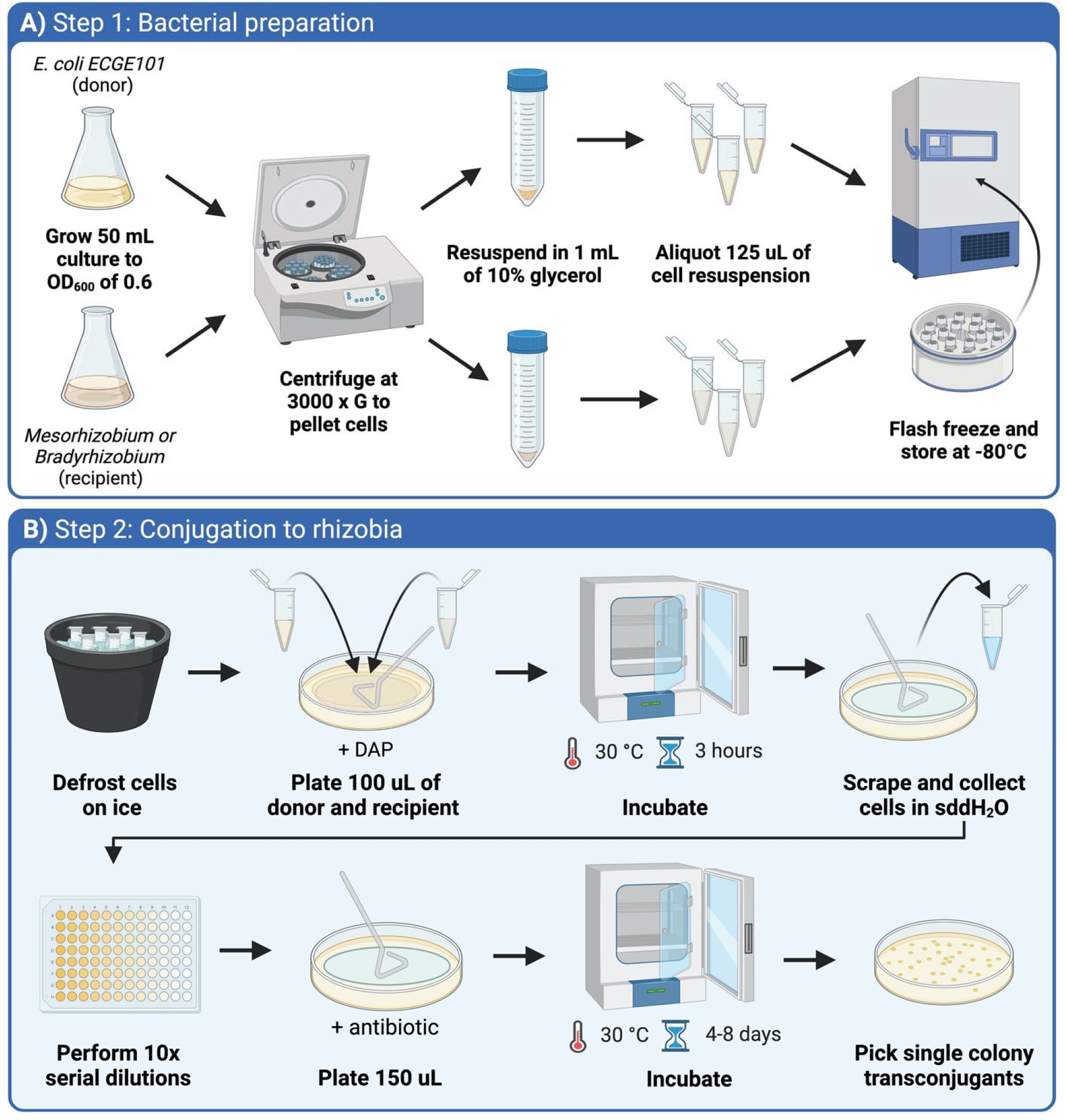
Schematic of the streamlined conjugation method for rhizobia. A) Outlines the preparation of the *E. coli* donor strain and rhizobial recipient strain. Both bacterial strains are grown to exponential phase before harvesting the cells. Pellets are resuspended in 10% glycerol and aliquoted into single conjugation reactions before being frozen for future use. B) Depicts steps for performing the conjugation. Frozen bacteria are defrosted and mixed on media supplemented with diaminopimelic acid (DAP) and incubated to allow for DNA transfer from donor to recipient. Cells are collected and diluted for transconjugant selection on media supplemented with appropriate antibiotics and lacking DAP.

### Conjugating shuttle vectors to rhizobia

Nine existing shuttle plasmids were investigated in this study; two contained only a broad-host-range origin of replication and the remaining seven also encoded a *repABC* operon from one of three endogenous rhizobial plasmids (Table S3). The RK2-based pFAJ1700 plasmid harbors the traditional incP vegetative origin, consisting of *oriV* and the *trfA* initiation protein, which can be induced to increase copy number from low to high (Doran, Konieczny and Helinski, 1998; Dombrecht, Vanderleyden and Michiels, 2001). Meanwhile, pTA-Mob2.0 uses the pBBR1 replication origin, that has medium copy number, but does not belong to common incompatibility groups (incP, incQ or incW) (Antoine and Locht, 1992; Jain and Srivastava, 2013; Soltysiak *et al*., 2019). These origins have been used in some rhizobia, but propagation is not consistent among all species. Other studies have demonstrated that multiple *repABC* plasmids can coexist in a cell without incompatibility at low copy number and origins from native rhizobial plasmids can be used in expression vectors for other species as well (Mazur *et al*., 2011; Żebracki *et al*., 2015). The pAGE/pBGE series of plasmids contain *repABC* operons from *S. meliloti* pSymA and pSymB, respectively, with three different antibiotic markers (*aadA* [spectinomycin], *tetA* [tetracycline] or *nptII* [neomycin and kanamycin]) (Brumwell *et al*., 2019). The third *repABC* operon, on pABCc-mob, originated from *Rhizobium etli* p42d and has the *nptII* antibiotic marker (Döhlemann *et al*., 2017). As such, it was hypothesized that at least one *repABC* shuttle plasmid would be able to replicate in the *Mesorhizobium* and *Bradyrhizobium* strains of interest.

Using the streamlined method, all nine plasmids were employed for conjugation to the three rhizobial strains. To create new conjugative donor strains, plasmid DNA was transferred into the DAP auxotroph with pTA-Mob by electroporation. Since pTA-Mob2.0 is a cis-acting conjugation plasmid, it did not require the presence of a helper plasmid (Soltysiak *et al*., 2019). Following DNA delivery, transconjugant colonies obtained on the lowest dilution were picked and purified by passing on selective media twice. Isolated DNA from each transconjugant was analyzed with multiplex PCR (MPX) to confirm the identity of the bacteria and presence of the shuttle plasmid of interest. Samples with amplicons that matched both positive controls - wildtype genomic DNA and plasmid DNA from the donor bacteria - demonstrated that the plasmid can be stably propagated in the recipient strain.

Two shuttle plasmids – pBGE1.0 and pABCc-mob – were stably replicated across all three strains and another – pBGE2.0 – was maintained in both *Mesorhizobium* strains. In *M. huakui*i NZP2235 (Figure 4) and *M. japonicum* R7A (Figure 5), four plasmids resulted in transconjugants that screened positive. For NZP2235, pAGE3.0 was also positive, showing maintenance of three different *repABC* operons. Meanwhile R7A was able to propagate the broad-host-range plasmid pFAJ1700, which is consistent with its use in previous studies (Ramsay *et al*., 2006; Sullivan, Brown and Ronson, 2013). For *Bradyrhizobium* sp. DOA9, there were just three shuttle plasmids that successfully propagated (Figure 6). Consistent with NZP2235, the third plasmid that screened positive was pAGE3.0, demonstrating replication of three *repABC* plasmids in this strain.

**Figure 4.**
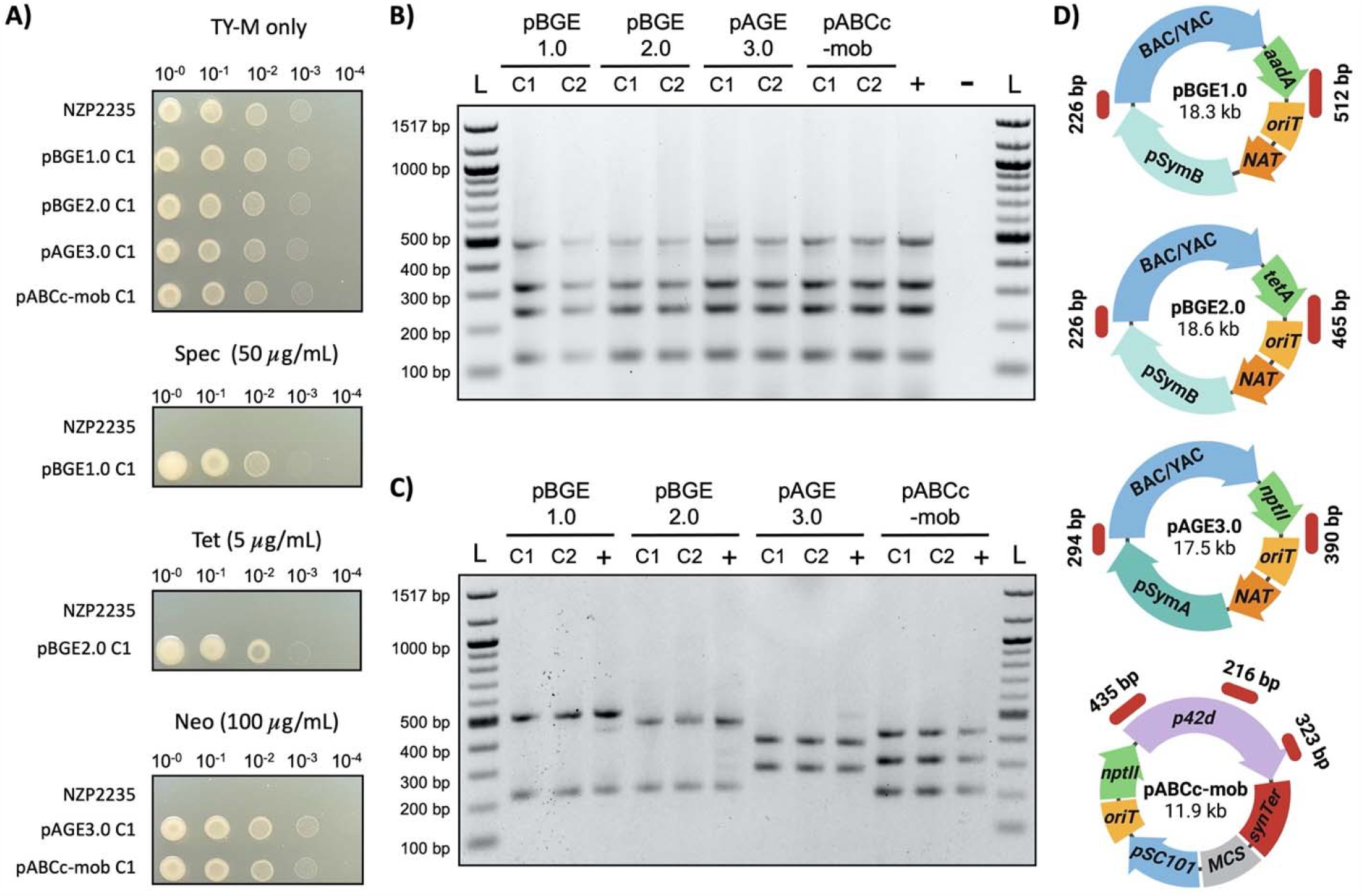
*M. huakuii* NZP2235 transconjugant colony analysis. A) Spot assays of one transconjugant colony on non-selective and selective media compared to the wildtype strain. B) Genomic DNA screened using multiplex PCR (MPX) for chromosomal marker regions (expected lengths: 132, 248, 325, and 502 bp). The positive control is DNA from wildtype NZP2235 and the negative control is no DNA template. C) Plasmid DNA screened using MPX with primers that amplify fragments indicated in the corresponding plasmid maps shown in panel D. The positive control is DNA isolated from the *E. coli* conjugative donor. D) Plasmid maps depicting the location and size of the expected amplicons for MPX screening (pBGE1.0 - 226 and 512 bp; pBGE2.0 - 226 and 465 bp; pAGE3.0 - 294 and 390 bp; pABCc-mob - 216, 323, 435 bp).

**Figure 5.**
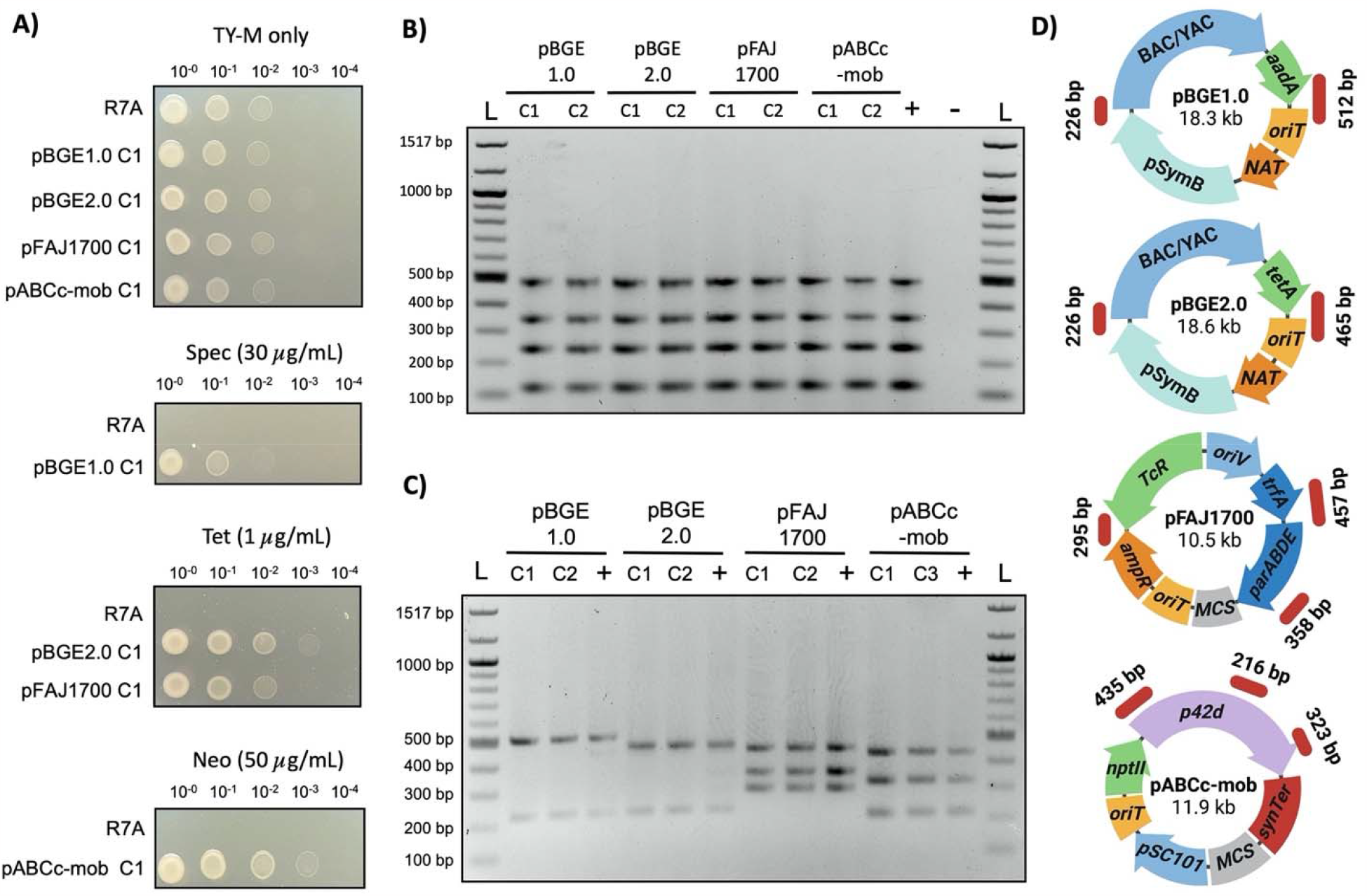
*M. japonicum* R7A transconjugant colony analysis. A) Spot assays of one transconjugant colony on non-selective and selective media compared to the wildtype strain. B) Genomic DNA screened using multiplex PCR (MPX) for chromosomal marker regions (expected lengths: 132, 248, 325, and 502 bp). The positive control is DNA from wildtype R7A and the negative control is no DNA template. C) Plasmid DNA screened using MPX with primers that amplify fragments indicated in corresponding plasmid maps shown in panel D. The positive control is DNA isolated from the *E. coli* conjugative donor. D) Plasmid maps depicting the location and size of the expected amplicons for MPX screening (pBGE1.0 - 226 and 512 bp; pBGE2.0 - 226 and 465 bp; pFAJ1700 - 295, 358, 457 bp; pABCc-mob - 216, 323, 435 bp).

**Figure 6.**
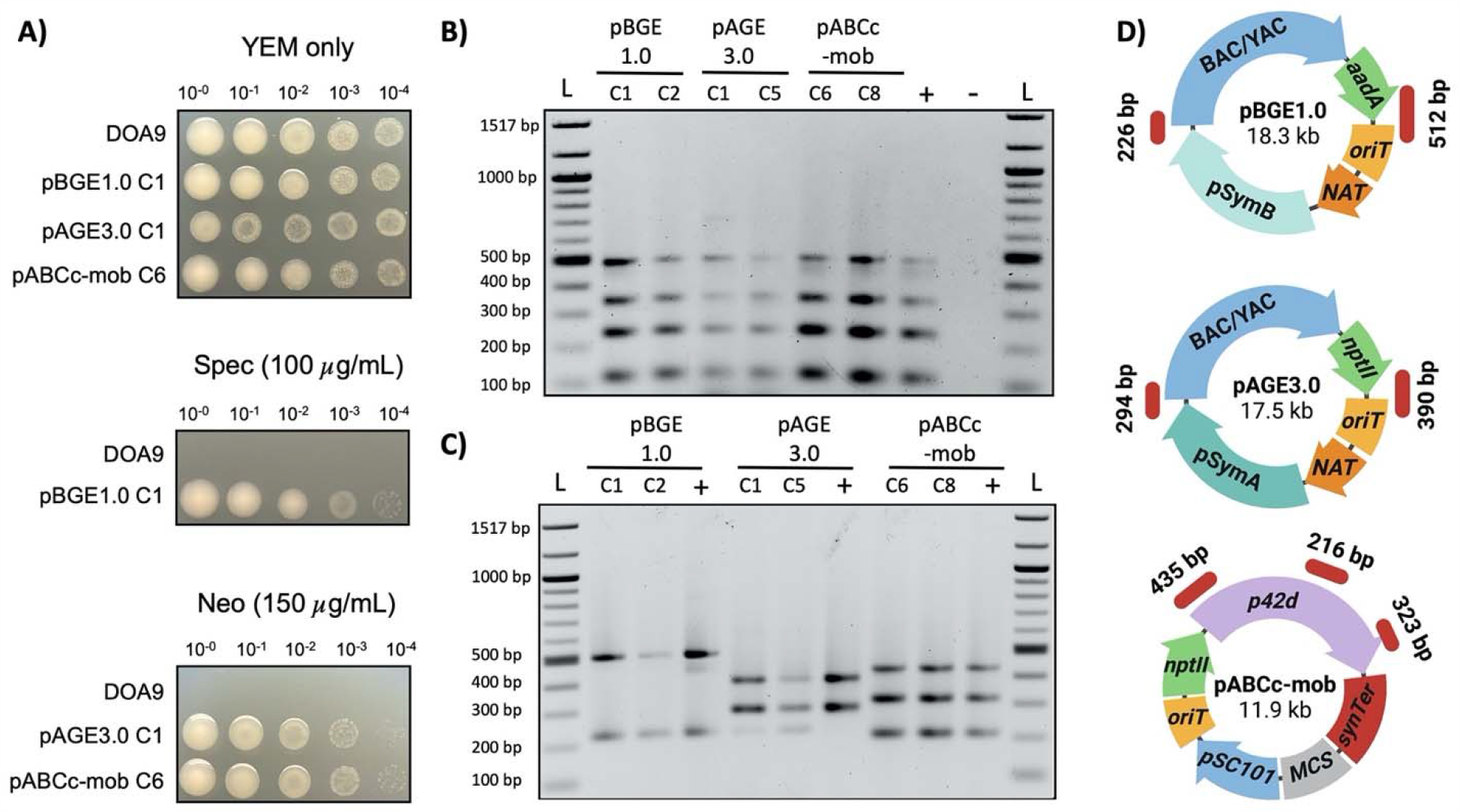
*Bradyrhizobium* sp. DOA9 transconjugant colony analysis. A) Spot assays of one transconjugant colony on non-selective and selective media compared to the wildtype strain. B) Genomic DNA screened using multiplex PCR (MPX) for chromosomal marker regions (expected lengths: 132, 248, 325, and 502 bp). The positive control is DNA from wildtype DOA9 and the negative control is no DNA template. C) Plasmid DNA screened using MPX with primers that amplify fragments indicated in corresponding plasmid maps shown in panel D. The positive control is DNA isolated from the *E. coli* conjugative donor. D) Plasmid maps depicting the location and size of the expected amplicons for MPX screening (pBGE1.0 - 226 and 512 bp; pAGE3.0 - 294 and 390 bp; pABCc-mob - 216, 323, 435 bp).

## CONCLUSION

In efforts to create a basic genetic toolbox for the three species of interest, their response to antibiotics was evaluated along with the propagation of a variety of shuttle plasmids following biparental mating. Given the limitations of working with wildtype bacteria that have fully intact R-M systems, a streamlined conjugation method was established for DNA delivery to any rhizobia species. Several plasmids were identified to be functional in the focal strains, with plasmids pBGE1.0 and pABCc-mob common among all three strains. Both plasmids were also previously shown to replicate in *S. meliloti* (Döhlemann *et al*., 2017; Brumwell *et al*., 2019). Given the successful use of pBGE1.0 and pABCc-mob across three rhizobial genera, it is likely that they will be of broad use across other phylogenetically diverse rhizobia. The availability of replicating plasmids in *M. japonicum* R7A, *M. huakuii* NZP2235, and *Bradyrhizobium* sp. DOA9 will greatly increase the genetic tractability of these organisms and can be customized for diverse purposes using a variety of cloning or assembly methods. The basic genetic toolbox developed in this work can be expanded by establishing promoter and terminator libraries for custom expression cassettes, gene editing via recombineering or CRISPR-Cas9 and large-scale whole genome engineering.

## ACKNOWLEDGEMENTS

We give thanks to Anke Becker’s laboratory of Philipps-Universitat Marburg, Germany for providing the pABCc-mob shuttle vector, Krzysztof Szczyglowski’s laboratory of Agriculture and Agri-Food Canada (AAFC), Canada for providing *M. huakuii* NZP2235 and Neung Teaumroong’s laboratory of Suranaree University of Technology, Thailand for *Bradyrhizobium* sp. DOA9. All figures in this publication were created using Biorender.com.

## AUTHOR CONTRIBUTIONS

JSM: conceptualization, formal analysis, investigation, methodology, writing – original draft, writing – review & editing; AKP: investigation; AJW: preliminary investigation; GCD: conceptualization, data curation, formal analysis, funding acquisition, methodology, resources, software, supervision, writing – original draft, writing – review & editing; BJK: conceptualization, formal analysis, funding acquisition, methodology, resources, supervision, writing – review & editing.

## DATA AVAILABILITY

Data generated or analyzed during this study are available in the published article and its supplementary materials, as well as at GenBank repository under accession numbers <u>CP139858</u> and <u>CP139859</u>.

## SUPPLEMENTAL MATERIALS

## REFERENCES

Antoine, R. and Locht, C. (1992) ‘Isolation and molecular characterization of a novel broadhost-range plasmid from Bordetella bronchiseptica with sequence similarities to plasmids from gram-positive organisms’, Molecular microbiology. Mol Microbiol, 6(13), pp. 1785–1799. doi: 10.1111/J.1365-2958.1992.TB01351.X.

Benmrid, B. et al. (2023) ‘Bioinoculants as a means of increasing crop tolerance to drought and phosphorus deficiency in legume-cereal intercropping systems’, Communications Biology 2023 6:1. Nature Publishing Group, 6(1), pp. 1–15. doi: 10.1038/s42003-023-05399-5.

Bochtler, M. (2021) ‘Distinction between self and non-self in restriction modification: The mysterious case of type IIL enzymes’, Structure. Cell Press, 29(6), pp. 512–514. doi: 10.1016/J.STR.2021.05.007.

Brumwell, S. L. et al. (2019) ‘Designer Sinorhizobium meliloti strains and multi-functional vectors enable direct inter-kingdom DNA transfer’. doi: 10.1371/journal.pone.0206781.

Burén, S. and Rubio, L. M. (2018) ‘State of the art in eukaryotic nitrogenase engineering’, FEMS Microbiology Letters. Oxford University Press, 365(2), p. 274. doi: 10.1093/femsle/fnx274.

Checcucci, A. et al. (2017) ‘Trade, diplomacy, and warfare: The Quest for elite rhizobia inoculant strains’, Frontiers in Microbiology. Frontiers Media S.A., 8, p. 312157. doi: 10.3389/fmicb.2017.02207.

Colombi, E. et al. (2021) ‘Comparative analysis of integrative and conjugative mobile genetic elements in the genus Mesorhizobium’, Microbial Genomics. Microbiology Society, 7(10), p. 657. doi: 10.1099/MGEN.0.000657.

Cournoyer, J. E. et al. (2022) ‘Engineering artificial photosynthetic life-forms through endosymbiosis’, Nature Communications. Nature Publishing Group, 13(1). doi: 10.1038/S41467-022-29961-7.

diCenzo, G. C. et al. (2018) ‘Robustness encoded across essential and accessory replicons of the ecologically versatile bacterium Sinorhizobium meliloti’, PLoS Genetics. Public Library of Science, 14(4). doi: 10.1371/journal.pgen.1007357.

diCenzo, G. C. et al. (2019) ‘Multidisciplinary approaches for studying rhizobium–legume symbioses’, Canadian Journal of Microbiology. Canadian Science Publishing, 65(1), pp. 1–33. doi: 10.1139/cjm-2018-0377.

diCenzo, G. C. et al. (2020) ‘Genome-scale metabolic reconstruction of the symbiosis between a leguminous plant and a nitrogen-fixing bacterium’, Nature Communications 2020 11:1. Nature Publishing Group, 11(1), pp. 1–11. doi: 10.1038/s41467-020-16484-2.

diCenzo, G. C. et al. (2023) ‘Refining the taxonomy of the order Hyphomicrobiales (Rhizobiales) based on whole genome comparisons of over 130 genus type strains’, bioRxiv. Cold Spring Harbor Laboratory, p. 2023.11.15.567303. doi: 10.1101/2023.11.15.567303.

diCenzo, G. C. and Finan, T. M. (2017) ‘The Divided Bacterial Genome: Structure, Function, and Evolution’, Microbiology and Molecular Biology Reviews. American Society for Microbiology, 81(3). doi: 10.1128/mmbr.00019-17.

Dixon, R. and Kahn, D. (2004) ‘Genetic regulation of biological nitrogen fixation’, Nature Reviews Microbiology 2004 2:8. Nature Publishing Group, 2(8), pp. 621–631. doi: 10.1038/nrmicro954.

Döhlemann, J. et al. (2017) ‘A Family of Single Copy repABC-Type Shuttle Vectors Stably Maintained in the Alpha-Proteobacterium Sinorhizobium meliloti’, ACS Synthetic Biology. American Chemical Society, 6(6), pp. 968–984. doi: 10.1021/acssynbio.6b00320.

Dombrecht, B., Vanderleyden, J. and Michiels, J. (2001) ‘Stable RK2-Derived Cloning Vectors for the Analysis of Gene Expression and Gene Function in Gram-Negative Bacteria’, Molecular Plant-Microbe Interactions MPMI, 14(3), pp. 426–430. doi: 10.1094/MPMI.2001.14.3.426.

Doran, K. S., Konieczny, I. and Helinski, D. R. (1998) ‘Replication Origin of the Broad Host Range Plasmid RK2’, Journal of Biological Chemistry. Elsevier BV, 273(14), pp. 8447–8453. doi: 10.1074/jbc.273.14.8447.

Ferri, L. et al. (2010) ‘Plasmid electroporation of Sinorhizobium strains: The role of the restriction gene hsdR in type strain Rm1021’, Plasmid. Plasmid, 63(3), pp. 128–135. doi: 10.1016/J.PLASMID.2010.01.001.

Geddes, B. A. et al. (2020) ‘The genomes of rhizobia’, Advances in Botanical Research. Academic Press Inc., 94, pp. 213–249. doi: 10.1016/bs.abr.2019.09.014.

Guo, K. et al. (2023) ‘Biological nitrogen fixation in cereal crops: Progress, strategies, and perspectives’, Plant Communications. Elsevier, 4(2), p. 100499. doi: 10.1016/J.XPLC.2022.100499.

Hawkins, J. P. and Oresnik, I. J. (2022) ‘The Rhizobium-Legume Symbiosis: Co-opting Successful Stress Management’, Frontiers in Plant Science. Frontiers Media S.A., 12, p. 796045. doi: 10.3389/fpls.2021.796045.

Jain, A. and Srivastava, P. (2013) ‘Broad host range plasmids’, FEMS Microbiology Letters. Oxford Academic, 348(2), pp. 87–96. doi: 10.1111/1574-6968.12241.

Jain, C. et al. (2018) ‘High throughput ANI analysis of 90K prokaryotic genomes reveals clear species boundaries’, Nature Communications 2018 9:1. Nature Publishing Group, 9(1), pp. 1–8. doi: 10.1038/s41467-018-07641-9.

Jarvis, B. D. W., Pankhurst, C. E. and Patel, J. J. (1982) ‘Rhizobium loti, a new species of legume root nodule bacteria’, International Journal of Systematic Bacteriology. Microbiology Society, 32(3), pp. 378–380. doi: 10.1099/00207713-32-3-378.

Kaneko, T. et al. (2000) ‘Complete genome structure of the nitrogen-fixing symbiotic bacterium Mesorhizobium loti’, DNA research□: an international journal for rapid publication of reports on genes and genomes. DNA Res, 7(6), pp. 331–338. doi: 10.1093/DNARES/7.6.331.

Karas, B. et al. (2005) ‘Invasion of Lotus japonicus root hairless 1 by Mesorhizobium loti Involves the Nodulation Factor-Dependent Induction of Root Hairs’, Plant Physiology. Oxford University Press, 137(4), p. 1331. doi: 10.1104/PP.104.057513.

Kaur, S. et al. (2023) ‘Complete Genome Sequences of the Species Type Strains Sinorhizobium garamanticum LMG 24692 and Sinorhizobium numidicum LMG 27395 and CIP 109850’, Microbiology Resource Announcements. American Society for Microbiology, 12(6). doi: 10.1128/mra.00251-23.

Kelly, S. et al. (2014) ‘Genome sequence of the Lotus spp. microsymbiont Mesorhizobium loti strain R7A’, Standards in Genomic Sciences. BioMed Central Ltd., 9(1), pp. 1–7. doi: 10.1186/1944-3277-9-6.

Krönauer, C. and Radutoiu, S. (2021) ‘Understanding Nod factor signalling paves the way for targeted engineering in legumes and non-legumes’, Current Opinion in Plant Biology. Elsevier Ltd, 62. doi: 10.1016/j.pbi.2021.102026.

Li, Q. and Chen, S. (2020) ‘Transfer of Nitrogen Fixation (nif) Genes to Non-diazotrophic Hosts’, Chembiochem□: a European journal of chemical biology. Chembiochem, 21(12), pp. 1717–1722. doi: 10.1002/CBIC.201900784.

Mazur, A. et al. (2011) ‘repABC-based replication systems of Rhizobium leguminosarum bv. trifolii TA1 plasmids: incompatibility and evolutionary analyses’, Plasmid. Plasmid, 66(2), pp. 53–66. doi: 10.1016/J.PLASMID.2011.04.002.

Meaney, R. S. et al. (2020) ‘Designer endosymbionts: Converting free-living bacteria into organelles’, Current Opinion in Systems Biology. Elsevier, 24, pp. 41–50. doi: 10.1016/J.COISB.2020.09.008.

Mehta, A. P. et al. (2018) ‘Engineering yeast endosymbionts as a step toward the evolution of mitochondria’, PNAS, 115(46). doi: 10.1073/pnas.1813143115.

Mehta, A. P. et al. (2019) ‘Toward a Synthetic Yeast Endosymbiont with a Minimal Genome’, Journal of the American Chemical Society. American Chemical Society, 141(35), pp. 13799–13802. doi: 10.1021/jacs.9b08290.

Meier-Kolthoff, J. P. et al. (2022) ‘TYGS and LPSN: a database tandem for fast and reliable genome-based classification and nomenclature of prokaryotes’, Nucleic Acids Research. Oxford Academic, 50(D1), pp. D801–D807. doi: 10.1093/NAR/GKAB902.

Meier-Kolthoff, J. P. and Göker, M. (2019) ‘TYGS is an automated high-throughput platform for state-of-the-art genome-based taxonomy’, Nature Communications 2019 10:1. Nature Publishing Group, 10(1), pp. 1–10. doi: 10.1038/s41467-019-10210-3.

Murray, N. E. (2000) ‘Type I Restriction Systems: Sophisticated Molecular Machines (a Legacy of Bertani and Weigle)’, Microbiology and Molecular Biology Reviews. American Society for Microbiology (ASM), 64(2), p. 412. doi: 10.1128/MMBR.64.2.412-434.2000.

Noisangiam, R. et al. (2012) ‘Genetic Diversity, Symbiotic Evolution, and Proposed Infection Process of Bradyrhizobium Strains Isolated from Root Nodules of Aeschynomene americana L. in Thailand’, Applied and Environmental Microbiology. American Society for Microbiology (ASM), 78(17), p. 6236. doi: 10.1128/AEM.00897-12.

Nora, L. C. et al. (2019) ‘Recent advances in plasmid-based tools for establishing novel microbial chassis’, Biotechnology Advances. Elsevier, 37(8), p. 107433. doi: 10.1016/J.BIOTECHADV.2019.107433.

Pankievicz, V. C. S. et al. (2019) ‘Are we there yet? The long walk towards the development of efficient symbiotic associations between nitrogen-fixing bacteria and non-leguminous crops’, BMC Biology 2019 17:1. BioMed Central, 17(1), pp. 1–17. doi: 10.1186/S12915-019-0710-0.

Ramsay, J. P. et al. (2006) ‘Excision and transfer of the Mesorhizobium loti R7A symbiosis island requires an integrase IntS, a novel recombination directionality factor RdfS, and a putative relaxase RlxS’, Molecular Microbiology. John Wiley & Sons, Ltd, 62(3), pp. 723–734. doi: 10.1111/J.1365-2958.2006.05396.X.

Riley, L. A. and Guss, A. M. (2021) ‘Approaches to genetic tool development for rapid domestication of non-model microorganisms’, Biotechnology for Biofuels 2021 14:1. BioMed Central, 14(1), pp. 1–17. doi: 10.1186/S13068-020-01872-Z.

Roer, L., Aarestrup, F. M. and Hasman, H. (2015) ‘The EcoKI Type I Restriction-Modification System in Escherichia coli Affects but Is Not an Absolute Barrier for Conjugation’, Journal of Bacteriology. American Society for Microbiology (ASM), 197(2), p. 337. doi: 10.1128/JB.02418-14.

Soltysiak, M. P. M. et al. (2019) ‘Trans-Kingdom Conjugation within Solid Media from Escherichia coli to Saccharomyces cerevisiae’, International Journal of Molecular Sciences. Multidisciplinary Digital Publishing Institute (MDPI), 20(20). doi: 10.3390/IJMS20205212.

Songwattana, P. et al. (2017) ‘Type 3 secretion system (T3SS) of Bradyrhizobium sp. DOA9 and its roles in legume symbiosis and rice endophytic association’, Frontiers in Microbiology. Frontiers Media S.A., 8(SEP), p. 287893. doi: 10.3389/FMICB.2017.01810.

Strand, T. A. et al. (2014) ‘A New and Improved Host-Independent Plasmid System for RK2-Based Conjugal Transfer’. doi: 10.1371/journal.pone.0090372.

Sullivan, J. T., Brown, S. D. and Ronson, C. W. (2013) ‘The NifA-RpoN Regulon of Mesorhizobium loti Strain R7A and Its Symbiotic Activation by a Novel LacI/GalR-Family Regulator’, PLOS ONE. Public Library of Science, 8(1), p. e53762. doi: 10.1371/JOURNAL.PONE.0053762.

Teamtisong, K. et al. (2014) ‘Divergent Nod-Containing Bradyrhizobium sp. DOA9 with a Megaplasmid and its Host Range’, Microbes and Environments, 29(4), pp. 370–376. doi: 10.1264/jsme2.ME14065.

Temme, K., Zhao, D. and Voigt, C. A. (2012) ‘Refactoring the nitrogen fixation gene cluster from Klebsiella oxytoca’, Proceedings of the National Academy of Sciences of the United States of America. National Academy of Sciences, 109(18), pp. 7085–7090. doi: doi.org/10.1073/pnas.1120788109.

Udvardi, M. and Poole, P. S. (2013) ‘Transport and metabolism in legume-rhizobia symbioses’, Annual review of plant biology. Annu Rev Plant Biol, 64, pp. 781–805. doi: 10.1146/ANNUREV-ARPLANT-050312-120235.

Waksman, G. (2019) ‘From conjugation to T4S systems in Gram-negative bacteria: a mechanistic biology perspective’, EMBO Reports. European Molecular Biology Organization, 20(2). doi: 10.15252/EMBR.201847012.

Walker, L., Lagunas, B. and Gifford, M. L. (2020) ‘Determinants of Host Range Specificity in Legume-Rhizobia Symbiosis’, Frontiers in Microbiology. Frontiers Media S.A., 11, p. 585749. doi: 10.3389/fmicb.2020.585749.

Wang, Q., Liu, J. and Zhu, H. (2018) ‘Genetic and molecular mechanisms underlying symbiotic specificity in legume-rhizobium interactions’, Frontiers in Plant Science. Frontiers Media S.A., 9, p. 313. doi: 10.3389/fpls.2018.00313.

Wang, S. et al. (2020) ‘Evolutionary Timeline and Genomic Plasticity Underlying the Lifestyle Diversity in Rhizobiales’, mSystems. American Society for Microbiology, 5(4). doi: 10.1128/msystems.00438-20.

Wongdee, J. et al. (2016) ‘nifDK Clusters Located on the Chromosome and Megaplasmid of Bradyrhizobium sp. Strain DOA9 Contribute Differently to Nitrogenase Activity During Symbiosis and Free-Living Growth’, Molecular Plant-Microbe Interactions. American Phytopathological Society, 29(10), pp. 767–773. doi: 10.1094/MPMI-07-16-0140-R.

Wongdee, J. et al. (2018) ‘Regulation of Nitrogen Fixation in Bradyrhizobium sp. Strain DOA9 Involves Two Distinct NifA Regulatory Proteins That Are Functionally Redundant During Symbiosis but Not During Free-Living Growth’, Frontiers in Microbiology. Frontiers Media S.A., 9(1644). doi: 10.3389/fmicb.2018.01644.

Wongdee, J. et al. (2023) ‘Role of two RpoN in Bradyrhizobium sp. strain DOA9 in symbiosis and free-living growth’, Frontiers in Microbiology. Frontiers Media S.A., 14, p. 1131860. doi: 10.3389/fmicb.2023.1131860.

Żebracki, K. et al./person-group>. (2015) ‘Plasmid-Encoded RepA Proteins Specifically Autorepress Individual repABC Operons in the Multipartite Rhizobium leguminosarum bv. trifolii Genome’, PLOS ONE. Edited by F. Hayes. Public Library of Science, 10(7), p. e0131907. doi: 10.1371/journal.pone.0131907.

