## Supplementary file for "Identifying functional multi-host shuttle plasmids to advance synthetic biology applications in *Mesorhizobium* and *Bradyrhizobium*"

METHODS

*Electroporation to* E. coli *ECGE101 to create conjugative donor strains*

*Competent cell preparation.* *E. coli* ECGE101 with pTA-Mob was streaked on LB agar supplemented with appropriate compounds and grown overnight at 37°C. A single colony was used to inoculate 5 mL of LB. The saturated overnight culture was diluted 1:100 (total 500 mL) and grown at 37°C to an OD_600_ of 0.6. The culture was chilled on ice for 30 minutes prior to centrifugation at 4000 x *g* for 15 minutes at 4°C. Supernatant was removed and the pellet was resuspended in cold sddH_2_O by gentle agitation in an ice bath. Centrifugation and washing were repeated twice with water and once with cold 10% glycerol. The cells were resuspended in a final volume of 1 mL of cold 10% glycerol. Aliquots of 100 µL were flash frozen in an ethanol bath and stored at -80°C.

*Electroporation.* Competent cells were thawed on ice for approximately 15 minutes before mixing 40 µL with 1.5 µL of DNA isolated from *E. coli.* The mixture was transferred to a pre-chilled 2 mm pathlength electrocuvette and pulsed with 2.5 kV using the BioRad Gene Pulser Xcell system. The cells were resuspended in 1 mL of SOC and recovered for one hour in a shaking incubator at 37°C. From the resuspension, 150 µL of cells were plated on selective LB medium and incubated at 37°C overnight. Single colonies were picked the following day.

*Alkaline Lysis for DNA Screening*

Cultures were centrifuged at 3,000 x *g* for 10 minutes and the supernatant was decanted. Residual media was collected and removed from the pellet. Cells were resuspended in 250 µL of P1 resuspension buffer (50 mM Tris·Cl, pH 8.0; 10 mM EDTA; 100 µg/mL RNase A) and transferred to a 1.5 mL Eppendorf tube. A 250 µL aliquot of P2 lysis buffer (200 mM NaOH. 1% SDS (w/v)) was added to the tube and inverted 6-8 times to mix. A 250 µL aliquot of P3 neutralization buffer (3.0 M potassium acetate, pH 5.5) was added and samples were inverted 6-8 times.  Tubes were centrifuged at 15,000 x *g* for 10 minutes. The supernatant was transferred to another 1.5 mL Eppendorf with 750 µL of ice-cold isopropanol and centrifugation was repeated. Supernatant was poured off the precipitated DNA and washed in 750 µL of ice-cold 95% ethanol. After the final centrifugation spin, the supernatant was removed and the pellet was allowed to dry before resuspending in 50 µL of TE buffer (10 mM Tris·Cl pH 8.0, 1 mM EDTA). Samples were incubated at 37^°^C to dissolve the DNA.

TABLES AND FIGURES

**Table S1.** Strains used and created in this study. Concentrations for antibiotic selection and supplement (supp.) are reported in µg/mL.

| Organism | Strain | Plasmid | Antibiotic | Supp. | Source |
| --- | --- | --- | --- | --- | --- |
| ***E. coli*** | DH5 | pABCc-mob | Kan100/Neo100 |  | Döhlemann (2017) |
|  |  | pFAJ1700 | Tet10/Amp100 |  | Dombrecht (2001) |
|  | Epi300 | pTA-Mob2.0 | Gm40 |  | Soltysiak (2019) |
|  | ECGE101 |  |  | DAP60* | Brumwell (2019) |
|  |  | pTA-Mob | Gm40 | DAP60 | Brumwell (2019) |
|  |  | pTA-Mob, pAGE1.0 | Gm40, Spec100 | DAP60 | Brumwell (2019) |
|  |  | pTA-Mob, pAGE2.0 | Gm40, Tet10 | DAP60 | Brumwell (2019) |
|  |  | pTA-Mob, pAGE3.0 | Gm40, Neo100 | DAP60 | Brumwell (2019) |
|  |  | pTA-Mob, pBGE1.0 | Gm40, Spec100 | DAP60 | Brumwell (2019) |
|  |  | pTA-Mob, pBGE2.0 | Gm40, Tet10 | DAP60 | Brumwell (2019) |
|  |  | pTA-Mob, pBGE3.0 | Gm40, Neo100 | DAP60 | Brumwell (2019) |
|  |  | pTA-Mob, pABCc-mob | Gm40, Kan100 | DAP60 | This study |
|  |  | pTA-Mob, pFAJ1700 | Gm40, Tet10 | DAP60 | This study |
|  |  | pTA-Mob2.0 | Gm40 | DAP60 | This study |
| ***M. huakuii*** | NZP2235 |  |  |  | Karas (2005) |
|  |  | pBGE1.0 | Spec50 |  | This study |
|  |  | pBGE2.0 | Tet5 |  | This study |
|  |  | pAGE3.0 | Neo100 |  | This study |
|  |  | pABCc-mob | Neo100 |  | This study |
| ***M. japonicum*** | R7A |  |  |  | Sullivan (2002) |
|  |  | pBGE1.0 | Spec30 |  | This study |
|  |  | pBGE2.0 | Tet1 |  | This study |
|  |  | pFAJ1700 | Tet1 |  | This study |
|  |  | pABCc-mob | Neo50 |  | This study |
| ***Bradyrhizobium* sp.** | DOA9 |  |  |  | Noisangiam (2012) |
|  |  | pBGE1.0 | Spec100 |  | This study |
|  |  | pAGE3.0 | Neo150 |  | This study |
|  |  | pABCc-mob | Spec100 |  | This study |

* Diaminopimelic acid (60 µg/mL)

**Table S2.** Putative genes that are predicted to be involved in R-M systems for the *Mesorhizobium* and *Bradyrhizobium* strains examined in this study.

| Organism | CDS | Genomic location (interval) | Length |
| --- | --- | --- | --- |
| *M. huakaii* NZP2235 | type I restriction-modification system subunit M* | Chromosome (4,099,296 <- 4,101,353) | 2058 bp (686 aa) |
|  | restriction endonuclease subunit S* | Chromosome (4,097,986 <- 4,099,299) | 1314 bp (438 aa) |
|  | type I restriction enzyme HsdR N-terminal domain-containing protein | Chromosome (5,273,970 <- 5,274,758) | 789 bp (263 aa) |
|  | Restriction endonuclease | Chromosome (1,251,948 -> 1,252,850) | 903 bp (301 aa) |
| *M. japonicum* R7A | Restriction endonuclease | Chromosome (215,912 <- 216,814) | 903 bp (301 aa) |
| *Bradyrhizobium* sp. DOA9 | type I restriction enzyme HsdR N-terminal domain-containing protein | Chromosome (4,994,191 -> 4,994,994) | 804 bp (268 aa) |

*****Genes belong to a single operon

**Table S3.** Plasmids used in this study and their characteristics.

| Plasmid | Size (kb) | Broad-host replication origin | *rep*ABC origin | Bacterial selection marker(s) | Eukaryotic selection marker(s) | Reference |
| --- | --- | --- | --- | --- | --- | --- |
| pTA-Mob | 52.7 | pBBR1 (*Bordetella bronchiseptica*) | N/A | Gm | N/A | Strand (2014) |
| pAGE1.0 | 18.1 | RK2 (*Pseudomonas aeruginosa*) | pSymA (*S. meliloti*) | Cm, Spc | HIS3, NAT | Brumwell (2019) |
| pAGE2.0 | 18.5 | RK2 (*Pseudomonas aeruginosa*) | pSymA (*S. meliloti*) | Cm, Tet | HIS3, NAT | Brumwell (2019) |
| pAGE3.0 | 17.5 | RK2 (*Pseudomonas aeruginosa*) | pSymA (*S. meliloti*) | Cm, Nm/Kan | HIS3, NAT | Brumwell (2019) |
| pBGE1.0 | 18.3 | RK2 (*Pseudomonas aeruginosa*) | pSymB (*S. meliloti*) | Cm, Spc | HIS3, NAT | Brumwell (2019) |
| pBGE2.0 | 18.6 | RK2 (*Pseudomonas aeruginosa*) | pSymB (*S. meliloti*) | Cm, Tet | HIS3, NAT | Brumwell (2019) |
| pBGE3.0 | 17.7 | RK2 (*Pseudomonas aeruginosa*) | pSymB (*S. meliloti*) | Cm, Nm/Kan | HIS3, NAT | Brumwell (2019) |
| pABCc-mob | 11.5 | pSC101 (*Salmonella typhimurium*) | p42d (*R. etli*) | Nm/Kan | N/A | Döhlemann (2018) |
| pFAJ1700 | 10.5 | RK2 (*Pseudomonas aeruginosa*) | N/A | Tet, Amp | N/A | Dombrecht (2001) |
| pTA-Mob2.0 | 56.5 | pBBR1 (*Bordetella bronchiseptica*) | N/A | Gm | HIS3 | Soltysiak (2019) |

**Table S4.** Primers used to screen bacteria with multiplex PCR. Genome primers were combined in a single mixture. Plasmid primers were grouped together according to the description. All oligonucleotides were synthesized by Integrated DNA Technologies (IDT).

| Name | Sequence (5’ to 3’) | | Description |
| --- | --- | --- | --- |
| Genome multiplex primers | | | |
| BK2623_F | GGTCAATTCGCGTTGATCGG | nifK gene (132 bp) | |
| BK2623_R | TCGGCCGAAAAAGTTCTCGA | nifK gene (132 bp) | |
| BK2624_F | ATCTACCAAGGGCCGGTACT | nifD gene (325 bp) | |
| BK2624_R | CTACAACATTGGCGGCGATG | nifD gene (325 bp) | |
| BK2625_F | TGCAAAAGCAATTCGGCGAA | nifA2 gene (502 bp) | |
| BK2625_R | TCCAGATTCCTGTTCGTGGC | nifA2 gene (502 bp) | |
| BK2627_F | CGAATAGCTGGCGAACTTGC | RpoN2 gene (248 bp) | |
| BK2627_R | GCTAGCGCTTCAAGATTGGC | RpoN2 gene (248 bp) | |
| Plasmid multiplex primers | | | |
| BK2147_F | GCGTGCATAATAAGCCCTACAC | pAGE1.0/pBGE1.0 (504 bp) | |
| BK2147_R | AGAGCAGGATTCCCGTTGAG | pAGE/pBGE antibiotic marker | |
| BK2148_F | GCCCTCTACTTGCTCTGCCTGC | pAGE2.0/pBGE2.0 (456 bp) | |
| BK2149_F | CTTTCTCTTTGCGCTTGCGT | pAGE3.0/pBGE3.0 (390 bp) | |
| BK2150_F | GGCGAGACATATGTGCCGTA | pAGE1.0/2.0/3.0 (294 bp) | |
| BK2150_R | CTCATCTGTCAGTGAGGGCC | pAGE/pBGE replication origin | |
| BK2151_F | AGTCGTTTCCGCAGTTGACA | pBGE1.0/2.0/3.0 (226 bp) | |
| BK2620_F | CTTCGATAACAGCAAGCGCC | pABCc-mob (216 bp) | |
| BK2620_R | CTGCGCGAGATGAAAGCTTC | pABCc-mob (216 bp) | |
| BK2621_F | ACATAGCCGCGATGACGATT | pABCc-mob (323 bp) | |
| BK2621_R | CGTTCAGGATCGCATGCAAG | pABCc-mob (323 bp) | |
| BK2622_F | GTCATGCGCCGTCCATAGTA | pABCc-mob (435 bp) | |
| BK2622_R | TCGCACCAGACCATATACGC | pABCc-mob (435 bp) | |
| BK2966_F | GCGGTATCATTGCAGCACTG | pFAJ1700 (295 bp) | |
| BK2966_R | TTTTCTACGGGGTCTGACGC | pFAJ1700 (295bp) | |
| BK2967_F | CAGGGGATCAAGATCGACGG | pFAJ1700 (457 bp) | |
| BK2967_R | TGATCTGCTGCTTCGTGTGT | pFAJ1700 (457 bp) | |
| BK2968_F | CCATTGCTGATGATCGGGGT | pFAJ1700 (358 bp) | |
| BK2968_R | AAGGTCGGGAAAATGCGCTA | pFAJ1700 (358 bp) | |
| BK3030_F | TTACATCGACATGCGCCTGG | pTA-Mob2.0 (421 bp) | |
| BK3030_R | TTGTTCTGTGCAGTTGGGTT | pTA-Mob2.0 (421 bp) | |
| BK3031_F | CATCCGCTTGCCGAATTCTG | pTA-Mob2.0 (194 bp) | |
| BK3031_R | TCTTTGGCATCGTCTCTCGC | pTA-Mob2.0 (194 bp) | |
| BK3032_F | GTCAATAAACCGGTAAACCAGCA | pTA-Mob2.0 (311 bp) | |
| BK3032_R | AGAGTCATCCGCTAGGTGGA | pTA-Mob2.0 (311 bp) | |

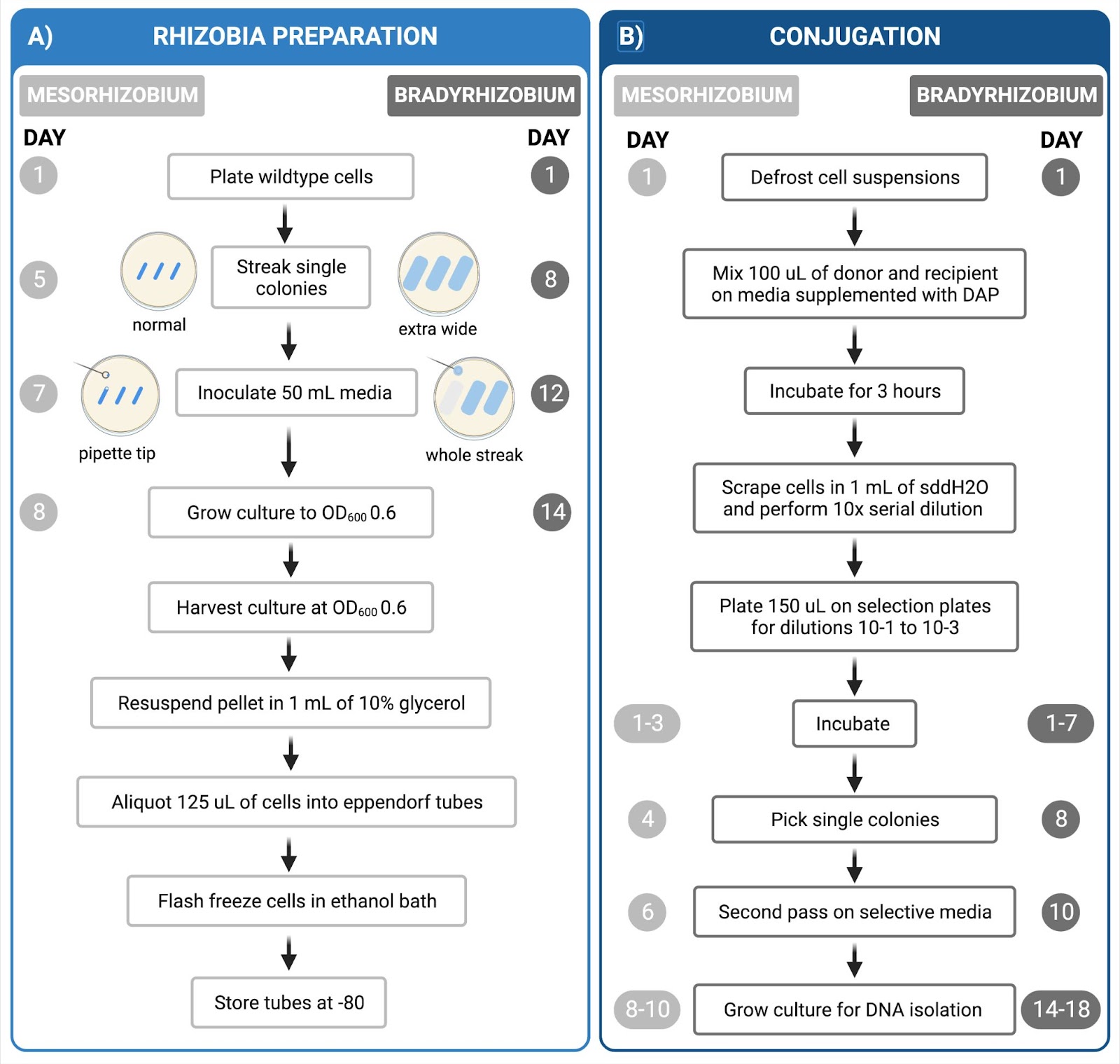

**Figure S1. Step-by-step protocol and time approximation for *Mesorhizobium* and *Bradyrhizobium* streamlined conjugation protocol.** A) Preparation of the rhizobial recipient takes eight days for intermediate-growing *Mesorhizobium*, and 14 days for slow-growing *Bradyrhizobium* from glycerol stock. *E. coli* donor preparation (not pictured) can be completed in four consecutive days. To inoculate the liquid cultures, a pipette-tip of cells was used for the *Mesorhizobium* strains while a patch (1x3 cm) of cells was used for the *Bradyrhizobium* strain. B) Thawed bacterial cells are used for conjugation. Single colonies are obtained after four or eight days for *Mesorhizobium* and *Bradyrhizobium*, respectively. Colonies were passed twice on selective agar media prior to growing culture for DNA isolation and screening.

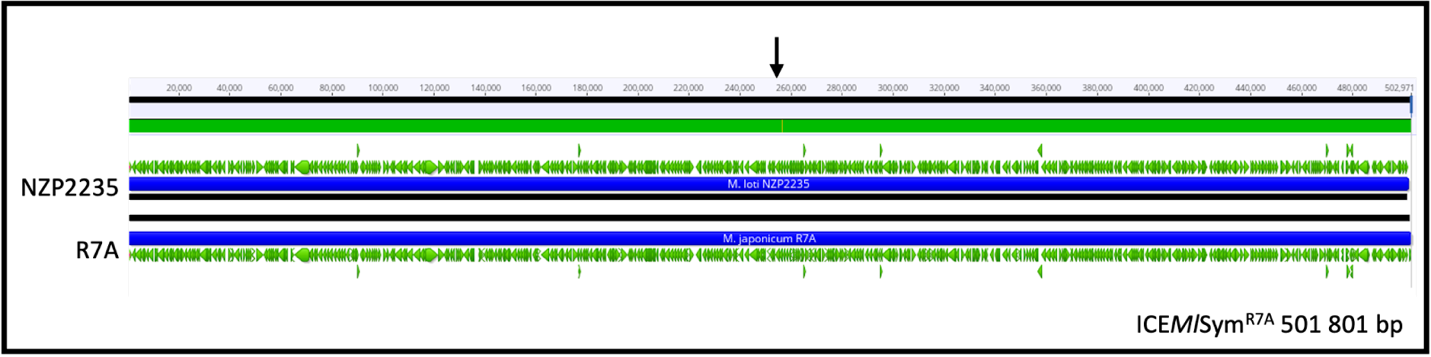

**Figure S2. Nucleotide sequence comparison of *M. japonicum* R7A and *M. huakuii* NZP2235 symbiotic island.** The 502 kb integrative and conjugative element (ICESym) from R7A was aligned to the NZP2235 chromosome using MAFFT *v7.490* on Geneious Prime 2023.2. Only a single nucleotide (indicated by the arrow) differentiated the sequences (99.9998% pairwise identity), resulting in an amino acid substitution in a Xaa-Pro peptidase family protein.

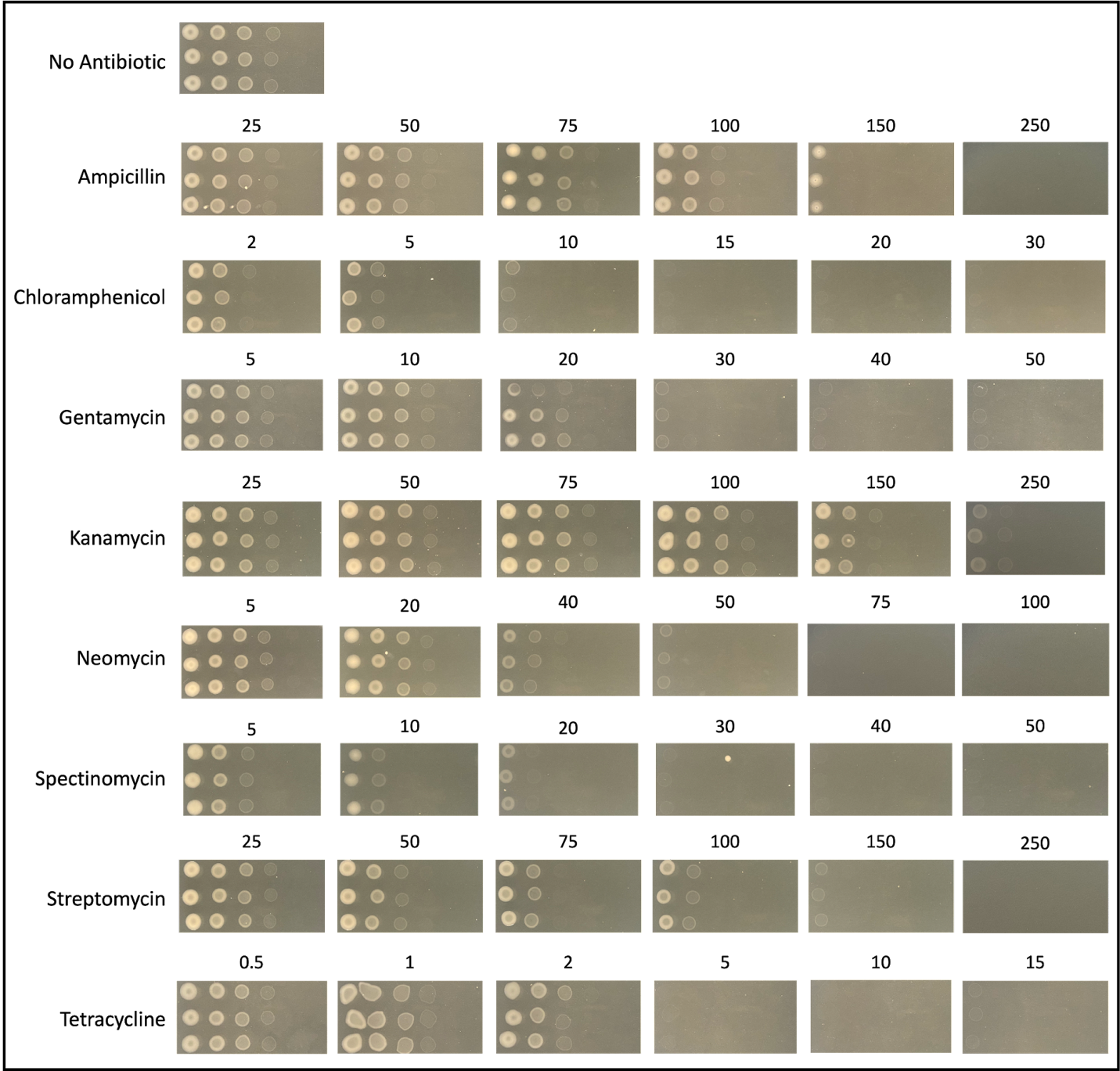

**Figure S3. Spot assay to assess natural antibiotic sensitivity of *M. huakuii* NZP2235.** Growth phenotype of wildtype cells exposed to various antibiotics and concentrations (µg/mL) on TY-M media. Single colony spotted in triplicate with 10-fold serial dilutions (10^-1^ to 10^-6^).

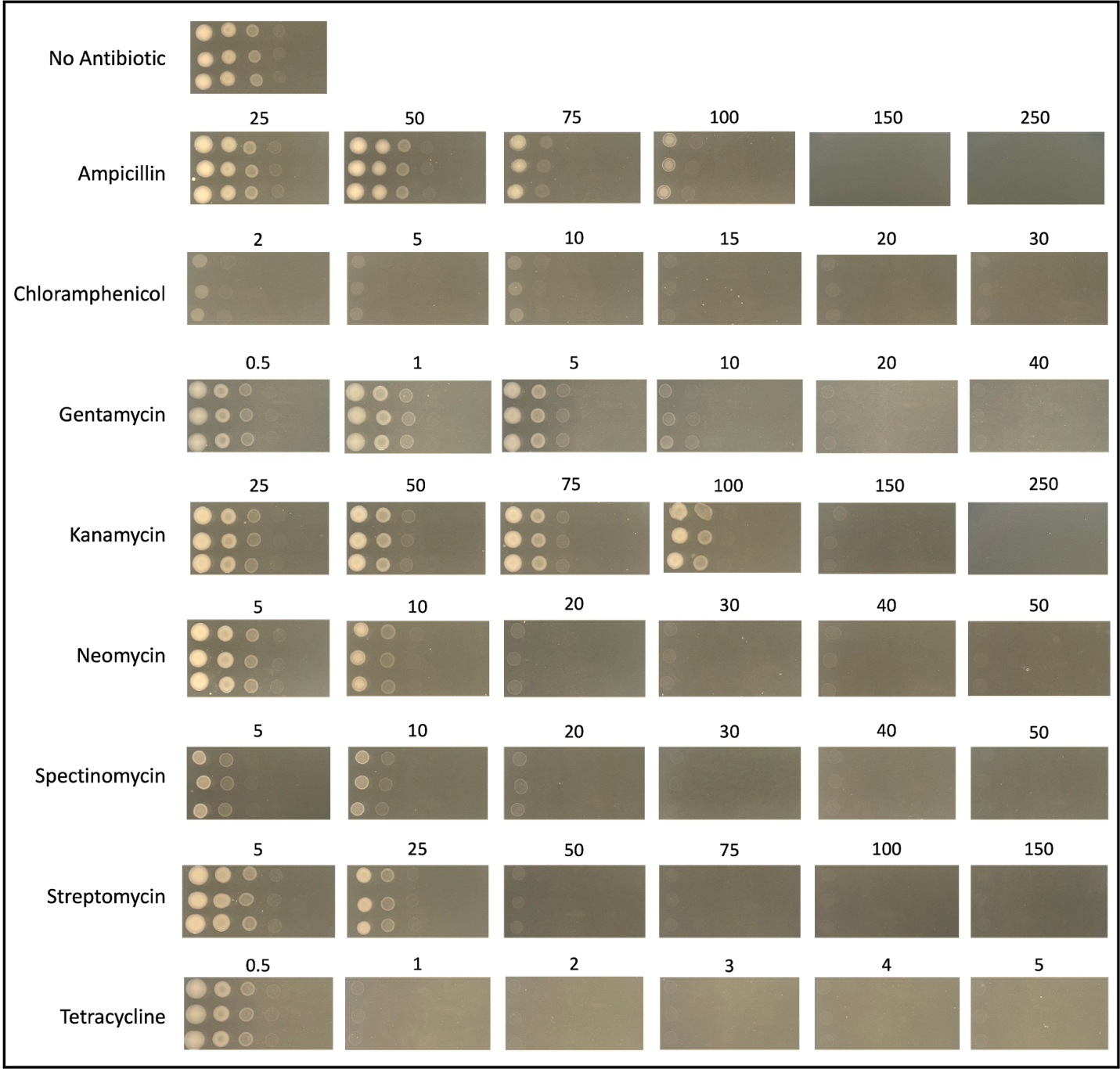

**Figure S4. Spot assay to assess natural antibiotic sensitivity of *M. japonicus* R7A.** Growth phenotype of wildtype cells exposed to various antibiotics and concentrations (µg/mL) on TY-M media. Single colony spotted in triplicate with 10-fold serial dilutions (10^-1^ to 10^-6^).

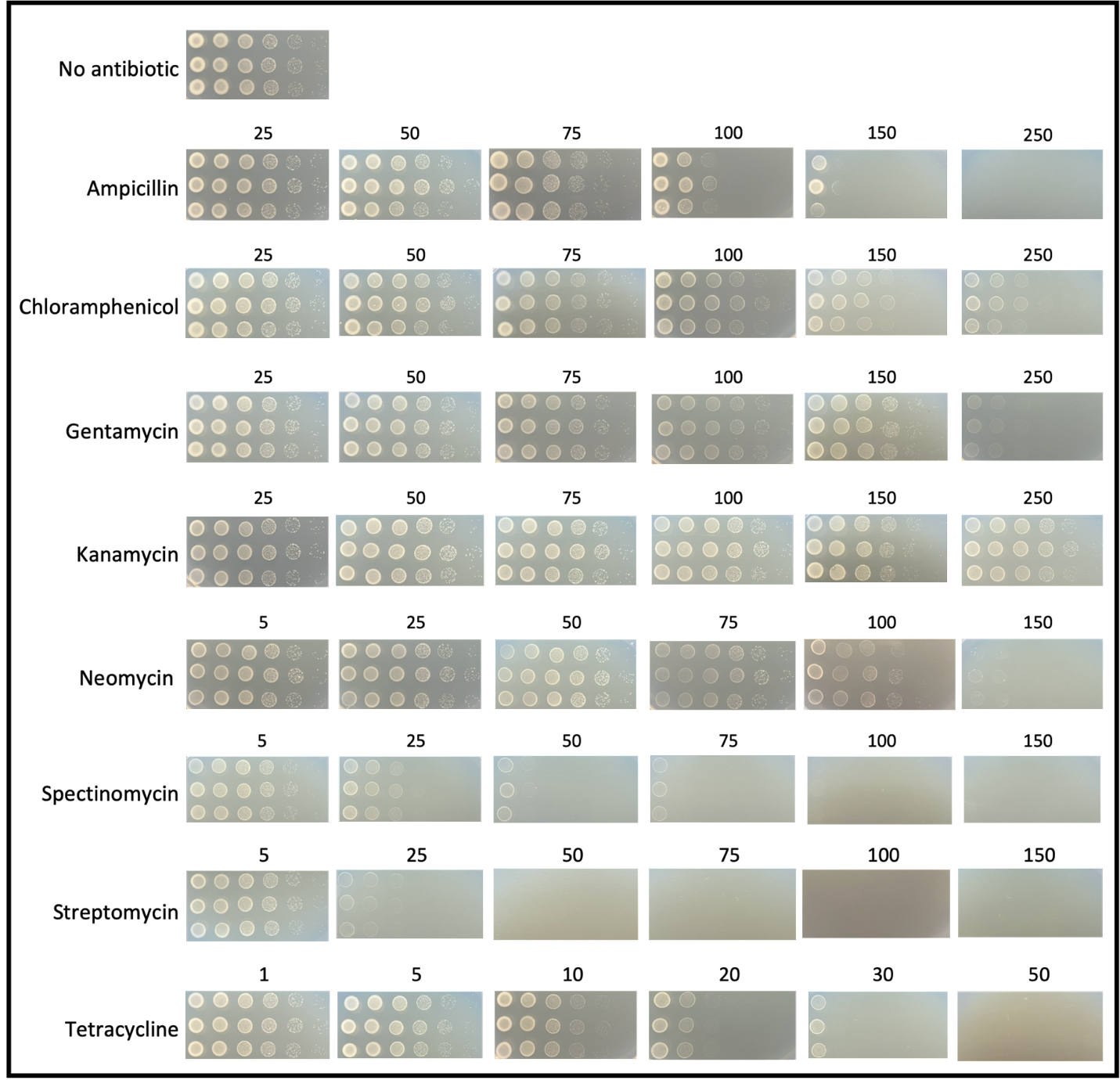

**Figure S5. Spot assay to assess natural antibiotic sensitivity of *Bradyrhizobium* sp. DOA9.** Growth phenotype of wildtype cells exposed to various antibiotics and concentrations (µg/mL) on YEM media. Single colony spotted in triplicate with 10-fold serial dilutions (10^-1^ to 10^-6^).
